# Differential Analysis Reveals Isoform Switching Following Pneumococcal Vaccination

**DOI:** 10.1101/2025.03.09.642237

**Authors:** Yeokyoung (Anne) Kil, Lior Pachter

## Abstract

Advances in RNA-sequencing (RNA-seq) technology have enabled scalable and accessible transcriptomics studies. Longitudinal RNA sequencing studies have been used to track gene expression over time, revealing biological pathways and expression patterns. Traditional approaches for such studies rely on pairwise comparisons or linear regression models, but these methods face challenges when dealing with many time points or modeling complex, non-linear expression patterns. Spline regression offers a robust alternative by efficiently capturing temporal patterns. In this study, we apply spline regression to analyze longitudinal RNA-seq data and demonstrate its advantages in isoform-level differential expression analysis. By modeling transcript-level expression, our method captures isoform switching events that can be obscured in traditional gene-level analyses.

## Introduction and Background

Advances in RNA-sequencing (RNA-seq) technology have made RNA-seq scalable and accessible (1), allowing researchers to conduct longitudinal transcriptomics studies that measure the expression of genes over time. Such studies have identified biological pathways and genes that display coordinated expression patterns (2–9) using a variety of methods that range from pairwise comparisons between time points to multivariate linear regression in time.

The most commonly used approaches for biological discovery from longitudinal RNA-seq studies are specific to the number of time points present in the study. Most published longitudinal transcriptomics studies, such as the aforementioned ones, have only 3-5 time points (2–8), making it difficult or impossible to apply traditional time series methods. In these cases where the number of time points is small, conducting pairwise comparisons between time points is the most intuitive and effective way to look for differentially-expressed genes. When there are more time points, pair-wise comparisons can still be useful for examining changes in gene expression at specific times, but the high-dimensionality of the analysis resulting from tens of thousands of genes and many time points can make implementation tedious, and aggregating results from the many pairs can be challenging and suffer from multiple testing difficulties.

One way to identify differentially expressed (DE) genes in time-course RNA sequences with many time points is to model the data with splines. A spline is a smooth function that consists of piecewise polynomials connected at points called “knots”, and fitting splines to time course data is a straightforward approach to reduce the dimensionality of the data, while preserving qualitative features. Splines can be fit to longitudinal data with uniformly spaced, or uneven time points in cases where sampling is not uniform. Moreover, spline regression can be applied to smooth non-linear data and is a convenient approach to modeling relationships between continuous variables and outcomes (10).

The shape of the polynomials that make up a spline and the number and placement of knots, define the many types of splines. The degree of the spline refers to the degree of the piecewise polynomials forming the spline: a cubic spline, for instance, is constructed from polynomials of degree 3. To ensure that a spline is continuous, there are two conditions that need to be met: 1) all polynomials of degree *d* in the spline must be (*d* − 1) times differentiable, and 2) all derivatives up to degree (*d* − 1) must be continuous at each knot. In addition to these conditions, a natural spline is constrained to be linear at the end points. Natural splines are widely used in spline regression because their edge constraints make them mimic natural phenomena more than other splines.

The number of knots in a spline can also be expressed in terms of “degrees of freedom,” which is a statistical term denoting the number of independently variable parameters in a system. The number of knots, *k*, and the degrees of freedom, *df*, are related to each other, but the relationship depends on the type of the spline. For example, B-splines, which are splines additionally defined by control points for improved shape control, have *df* = *k* + *d* degrees of freedom, where *d* denotes the degree of the polynomials (11). Natural splines have *df* = *k* + 1 degrees of freedom, due to the constraints at the end points (11). The number and placement of knots can determine how well the spline fits the data, so it is important to choose an appropriate number of knots for the spline. With more knots the spline fit may be better, but additional parameters increase the risk of overfitting (10). As for the location of the knots, they can be placed uniformly throughout the range of the data with equal numbers of points between each knot, or be optimized to minimize variance (12).

An important consideration in analyzing RNA-seq data is the concept of gene isoforms—different RNA transcript variants produced from the same gene. These variants result from processes like alternative splicing, where different exons can be included or excluded from the final transcript (13). Isoforms from the same gene can encode mRNA or protein-swith different functions, localizations, or regulatory properties (13). A particularly significant phenomenon is isoform switching, where the relative abundance of these transcript variants changes between conditions or over time (14). This switching can reveal subtle but biologically important signals that would be completely masked in conventional gene-level analyses where transcript counts are simply summed together (15). Detecting these isoform-level changes is especially valuable in longitudinal studies of complex biological processes, such as immune responses, where the specific mRNA isoform being expressed can dramatically affect biological outcomes.

In this work, we analyze a dataset published in Mias et. al. (16), which aimed to demonstrate the potential of saliva multiomics for non-invasive health monitoring by tracking immune responses to a vaccination (16). Researchers monitored one healthy individual across three distinct time periods: (1) hourly sampling for 24 hours under normal conditions, (2) hourly sampling for 24 hours after administering the pneumococcal polysaccharide vaccine (PPSV23), and (3) daily sampling for 33 days following the vaccination. The original study utilized a pipeline named MathIOmica, which employs spectral methods to analyze time series data based on temporal trends through autocorrelation analysis, where each time series is compared with a delayed version of itself and categorized into classes based on autocorrelations and signal spikes (16). From the original data, we sub-sample and construct two time series, hourly and daily, from the bulk RNA-seq data generated from this study and show that spline regression coupled with accurate isoform-level quantification can be used to identify isoforms that may be biologically relevant, but that are missed in a naïve gene-level analysis.

## Results

### Description of the Time Series

The sub-sampled hourly series has 20 time points prior to vaccination and 20 time points after vaccination. The pre-vaccination hourly time points are matched with the post-vaccination time points for the hour of the day, and time points are evenly distributed. The principal components analysis (PCA) of the hourly samples shows some separation between early and late time points, but not much separation between conditions (Figure 2, left). The sub-sampled daily series has 30 time points, all of which are post-vaccination, sampled at the same time every day. The PCA of the daily samples did not show a conclusive pattern on the time-course behavior of the samples (Figure 2, right).

**Fig. 1.**
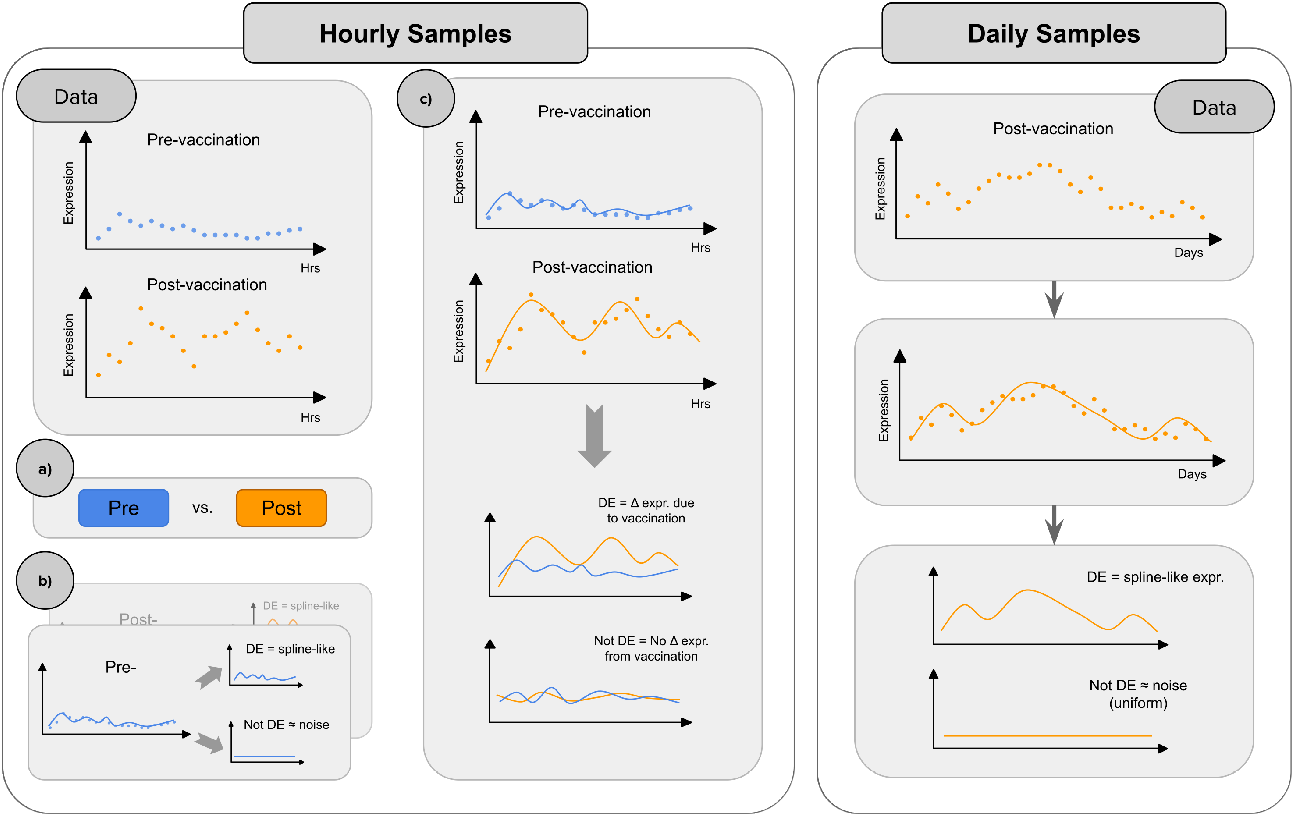
Data and analysis workflow. Left: Hourly samples with 20 pre- and 20 post-vaccination time points, matched by hour. a) Broad DE analysis between pre- and post-vaccination without time consideration. b) DE analysis within each condition to detect temporal expression changes. c) DE analysis with both time and condition, comparing splines fit to each. Right: Daily samples with 30 post-vaccination time points. Splines are fit to each isoform, and DE isoforms with spline-like patterns are identified.

**Fig. 2.**
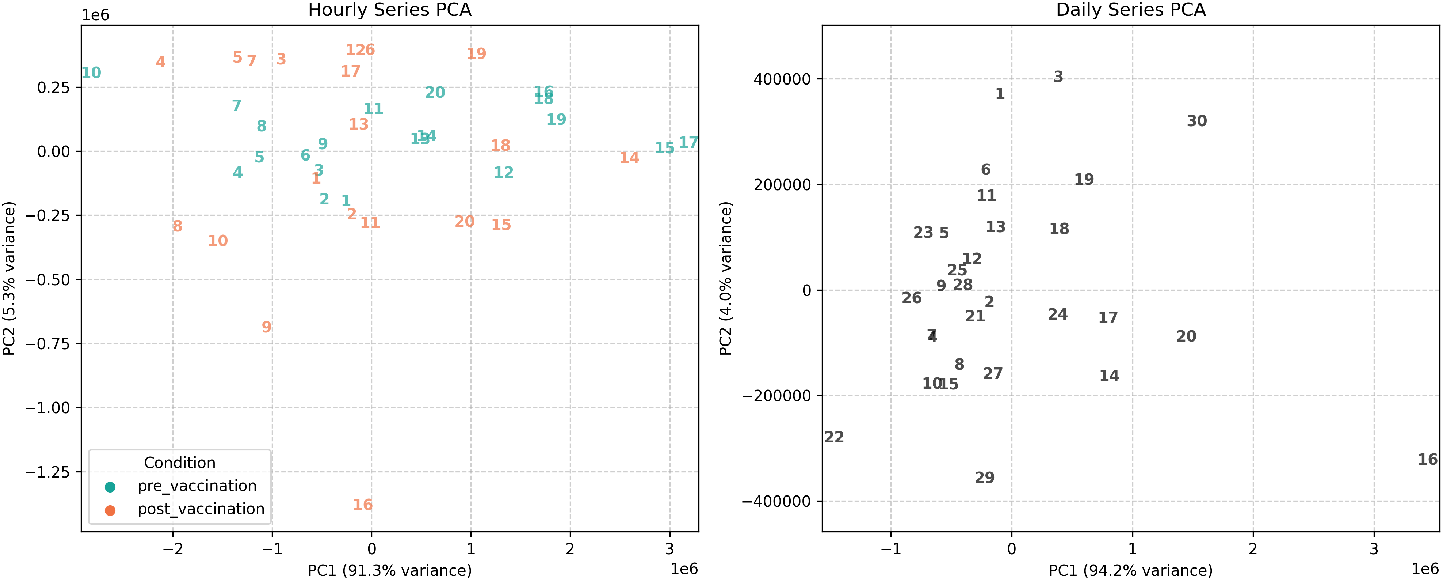
PCA of hourly (left) and daily (right) samples, colored by condition and labeled by time. Left) For the hourly series, there is some separation between the early and late time points but not much separation between pre- and post-vaccination samples. Right) Overall, the samples are more clustered than spread out, and the PCA does not show a conclusive pattern on how samples change over time.

### Differential Gene Expression from Immune Perturbation

#### Hourly Series, Pre- and Post-vaccination

From the hourly series with 20 pre- and 20 post-vaccination samples, a total of 1208 genes were identified as differentially expressed over time and between conditions (Supplementary Result 1). The DE genes from the hourly series are presented in Supplementary Result 1, ranked by statistical significance. Among the top differentially-expressed (DE) genes are TYROBP, STXBP2, PCBP2, and ADGRE5, which are genes associated with immune system related pathways, such as the Immune System pathway (R-HSA-168256) and Innate Immune System pathway (R-HSA-168249) from the Reactome 2024 database (17). TYROBP, one of the top DE genes from the hourly series, is a key immune signaling adaptor that regulates the activation of natural killer cells and myeloid cells, playing a crucial role in innate immunity and neuroinflammation (18, 19). 178 of the 1208 DE genes from the hourly series were also found in the original study (16), where many of them were grouped into the Lag 1 cluster, known for providing key information about time series changes (Supplementary Figure 1). The relatively small overlap between genes identified in the original study and our results may be related to the fact that the original study subtracted pre-vaccination counts from post-vaccination counts prior to differential expression analysis, whereas we compared the two time series as they are, which could account for the differences in identified genes.

Gene Set Enrichment Analysis (GSEA) for Reactome pathways using all 1208 DE genes showed numerous pathways expected of an immune response to a vaccination, such as “Immune System,” “Innate Immune System,” and “Neutrophil Degranulation,” but there were pathways not specific to immune response, such as “Cellular Responses to Stimuli/Stress” (Supplementary Result 2). In spite of the differences in Reactome databases used for GSEA between the original study and our results, terms from the Lag 1 cluster of the original study (Reactome 2022) and DE genes from sleuth (Reactome 2024) showed significant overlap in immune-system-related terms, suggesting that our spline-based analysis was able to capture the immune response as well. Overlapping GSEA terms included “Adaptive Immune System,” “Immune System,” “Innate Immune System,” and “Neutrophil Degranulation,” but also many non-immune-related pathways.

#### Daily Series, with one condition

In the daily series with 30 post-vaccination samples, sleuth identified 241 genes with expression patterns that varied over time more than noise as differentially expressed genes (Supplementary Result 1). The DE genes from the daily series are presented in Supplementary Result 1, ranked by statistical significance. PLAT, the top DE gene from the daily series, is known to modulate both innate and adaptive immunity by influencing immune cell activation, migration, and cytokine production (20). 123 out of the 241 DE genes were also marked as DE in the original study, with most belonging to the SpikeMin cluster, and the remainder primarily in the Lag 1 cluster (Supplementary Figure 2). The substantial overlap with the SpikeMin cluster suggests that spline regression with sleuth identified many genes with abnormally low levels at specific points in the time course.

GSEA for Reactome pathways with DE genes from the daily series revealed many significant pathways related to immune responses, such as “Immune System,” “Neutrophil Degranulation,” and “Cytokine Signaling in Immune System” (Supplementary Result 2). However, similar to the hourly series, the top pathways contained Reactome pathways that are not specific to immune response, such as “Metabolism of Proteins.” As with the hourly series, GSEA terms from the Lag 1 cluster of the original study (Reactome 2022) and DE genes from sleuth (Reactome 2024) showed overlap in immune-system-related terms, such as “Adaptive Immune System,” “Immune System,” “Innate Immune System,” “Interferon Gamma Signaling,” and “Neutrophil Degranulation,” showing that our methods were able to capture pathways similar to the original study that are relevant to the immune response.

### Detection of Isoform Switching through Isoform-Resolution Analysis

Using kallisto(21) and sleuth(22) in tandem enables differential gene expression analysis on the gene isoform resolution. With spline regression on each isoform, we can capture isoform-level temporal dynamics that could be hidden if isoform counts are aggregated into gene-level counts prior to analysis. For example, if two transcripts have opposing temporal expression patterns, adding them together for a gene-level analysis may cancel out their expression dynamics. We show that performing differential expression analysis in the isoform level captures genes more specific to the immune response. Furthermore, we show that for many of the DE genes in both the hourly and daily series, isoform-level results show isoform switching, which is defined as a significant change in the relative contribution of isoforms to the expression of the parent gene (14).

#### Hourly Series

We ran sleuth in transcript mode with p-value aggregation enabled on abundance estimates from spline regression applied to each isoform. The gene-level DE results were compiled by aggregating p-values of the isoforms with the Lancaster method. As sleuth allows for gene-level analysis, we analyzed the hourly series on both isoform-level and gene-level and compared the results. 615 genes were identified as differentially expressed from gene-level analysis. Of the 615 genes, 255 were commonly identified as DE genes in both gene-level and isoform-level analyses (Supplementary Figure 3). The top DE gene for both gene- and isoform-level analyses was TMPRSS11E, but the overlap between the two analyses was small otherwise. The vast difference in DE genes from gene-level and isoform-level analyses may come from how sleuth aggregates the isoforms for each of the analyses: in gene-mode, sleuth aggregates all isoforms for a given gene prior to filtering and analysis, which may lead to isoforms with highly contrasting patterns getting added to form a lower contrasting pattern and getting filtered out as a result. GSEA with the 615 DE genes from gene-level analysis identified functional pathways not particularly relevant to the immune response (Supplementary Table 1), whereas GSEA with DE genes from isoform-level analysis showed immune-related pathways, as expected from a longitudinal study with an immune perturbation. Together, these results demonstrate that isoform-level analysis is not only more sensitive in detecting differential expression, but also more effective at capturing biologically relevant signals alligned with the known immune response.

When running sleuth directly on isoform-level abundance estimates, we were able to capture genes with isoforms that show varying expression patterns over time. One of the top genes, VASP, showed evidence of isoform switching, as shown in Figure 3. The post vaccination splines for all 8 isoforms of VASP are plotted in the top panel, as well as a breakdown of each isoform’s expression (bottom left) compared to the aggregate gene-level expression (bottom right). The aggregated gene-level expression over time looks different from most of the splines, suggesting that dynamics of temporal expression are not the same for all isoforms and that summing the isoform counts may hide individual isoform expression patterns. We can see from the isoform fractions that the isoform that dominates the gene’s expression switches from ENST00000588273 to ENST00000705987, and later to ENST00000245932. This isoform switching phenomenon in VASP is particularly interesting in this time series with immune perturbation, as VASP is known to play a crucial role in T-cell activation and expansion (23). Although results are visualized for VASP as an example, isoform switching can be observed in many of the DE genes. However, this phenomenon was not limited to post-vaccination and was also observed in pre-vaccination series, despite the fact that pre-vaccination splines tended to be more “flat” compared to post-vaccination splines for a given isoform.

**Fig. 3.**
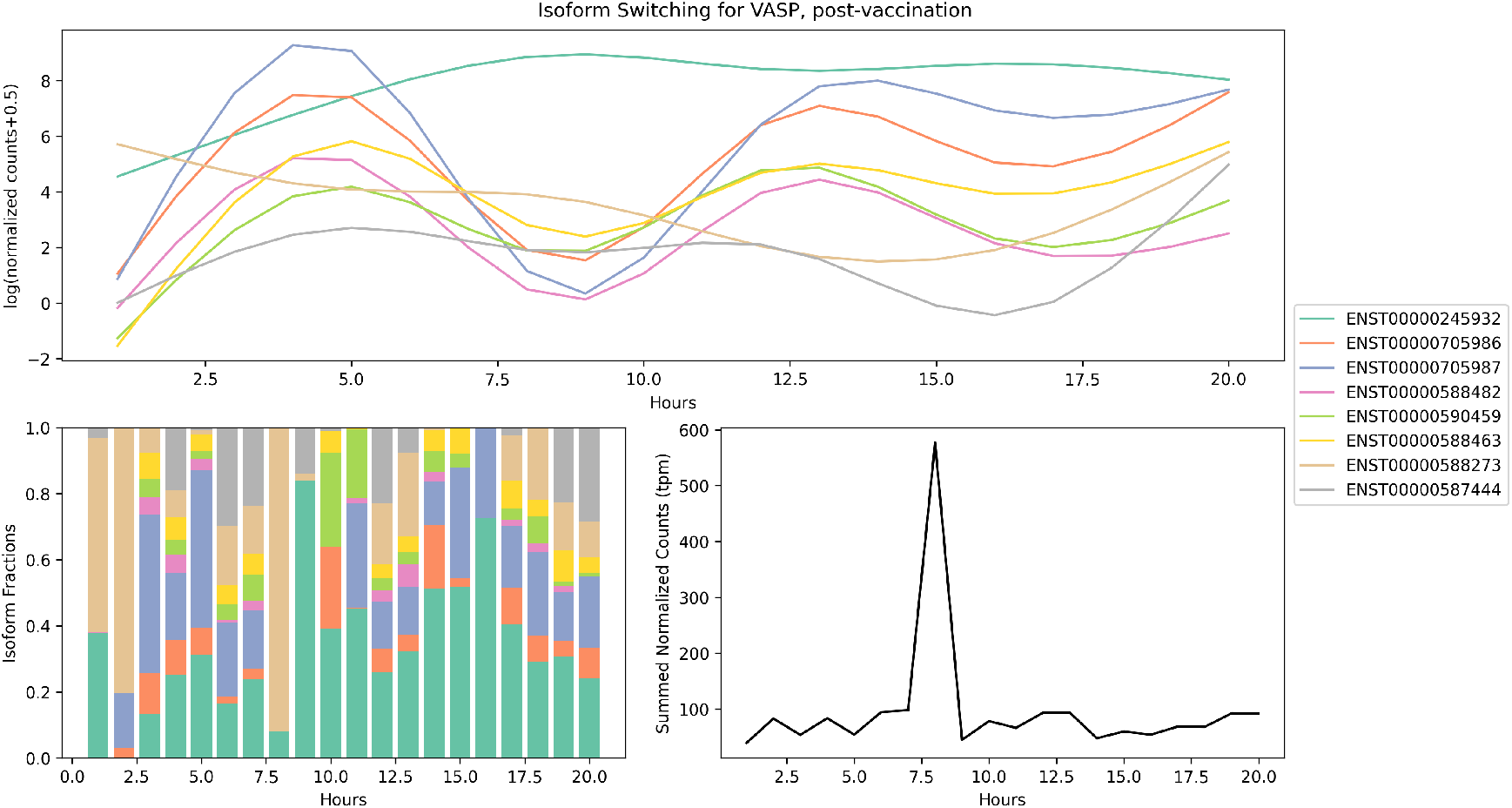
Isoform switching seen in gene VASP, in post-vaccination series. Top: Overlayed isoform expression patterns show different temporal trends. Bottom left: Fraction of isoform expression of total gene expression over time, showing change in which isoform dominates the expression of the gene. Bottom right: Total gene expression over time, which can deviate from the constituent isoform expression patterns, implying that isoform-level analysis may highlight temporal dynamics of a gene that might be missed on a gene-level analysis.

#### Daily Series

The difference between gene-level and isoform-level analysis results were also observed in the daily series. While 241 genes were marked as DE in the isoform-level analysis with p-value aggregation, only 106 genes were marked as DE in the gene-level analysis. Out of the 241 genes identified as DE from the isoform-level analysis, only 29 were also identified as DE on the gene level (Supplementary Figure 4). The top 10 DE genes from isoform-level analysis, except for RPS27L, were not identified as DE in gene-level analysis. As with the hourly series, GSEA with the 106 genes from gene-level analysis identified functional pathways that were not specific to an immune response that is expected of a vaccination (Supplementary Table 2), suggesting that many of the genes that were relevant to the immune response were lost in gene-level analysis and that isoform-level analysis was more sensitive to differentially expressed genes, especially those that were biologically relevant.

Similar to the hourly series, DE analysis on the isoform level identified genes with varying isoform expression patterns, with some showing signs of isoform switching. Figure 4 shows the temporal expression of isoforms for gene PLAT, the top DE gene from the daily series, as an example. For the five isoforms that passed the filter for low expression and were aggregated into the PLAT gene, there seem to be distinct groups of splines that resemble each other (Fig. 4, top panel). This is further corroborated by the isoform fractions (Fig. 4, lower left) as it shows ENST00000352041 and ENST00000677722 taking up most of the expression of PLAT in the first few days of the series and then later being replaced by ENST00000429089 and ENST0000067867 later in the series. While this switch in isoform fractions can also be observed in other DE genes from the isoform-level results, it does not seem to be universal among all DE genes, especially in those where all isoforms are consistently expressed over time.

**Fig. 4.**
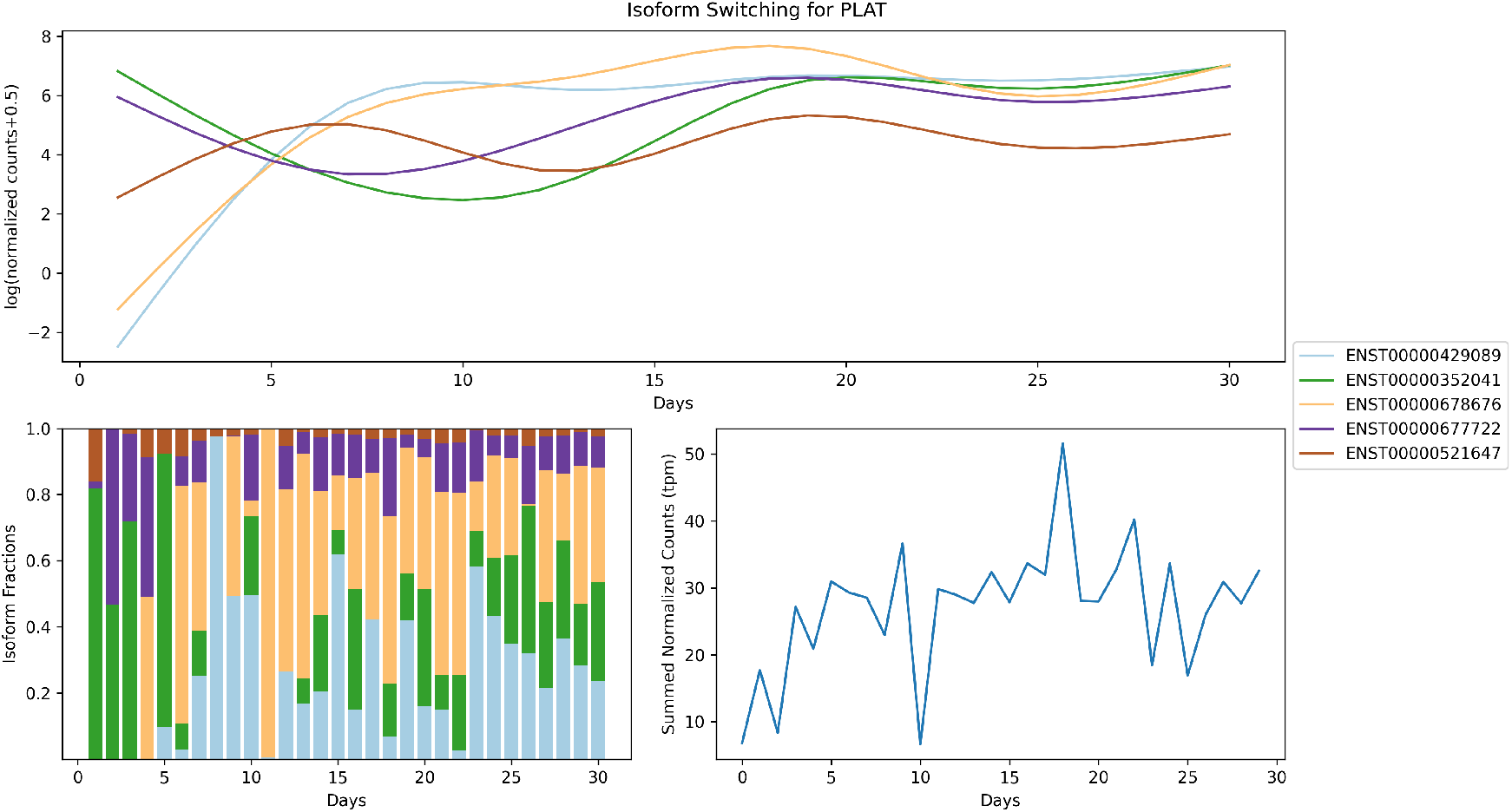
Isoform switching seen in gene PLAT from the daily time series. Top: Overlayed isoform expression patterns show different temporal dynamics of isoforms for PLAT. Bottom left: Fraction of isoform expression to total gene expression shows the change in which isoform dominates the expression for the gene. Bottom right: Total expression for gene PLAT, which does reflect the complexity of isoform expression patterns as evident in the hourly series.

## Discussion

We present a method of analyzing longitudinal data that leverages sleuth for time-course DE analysis at the isoform-level using spline regression, integrating both time and condition as covariates. Spline regression is shown to be an effective method of modeling non-linear, complex temporal gene expression, compared to other methods of analyzing longitudinal data. Furthermore, sleuth’s ability to perform isoform-level DE analysis enables the identification of genes with isoform switching that may be missed in a classic, gene-level DE analysis. The findings from this study demonstrate the potential of this approach for more precise and comprehensive analysis of time-dependent transcriptomic data in similar longitudinal studies with multiple experimental conditions and many time points.

The concept of analyzing time-course RNA-seq with splines is not new. The analysis has been introduced as an approach that can be utilized with different DE analysis tools vsuch as edgeR (24) and limma (25). The novelty of our approach is to apply spline regression on the isoform level and to aggregate them into gene-level conclusions via p-value aggregation with sleuth. To contextualize our results, we bench-marked our method against an edgeR-limma-based approach using transcript-level counts, following their recommended procedures (25). Despite using similar spline models, edgeR-limma identified only 10 differentially expressed genes from the hourly series, whereas our approach detected significantly more. This difference highlights the sensitivity of our p-value aggregation method (Supplementary Result 3).

Splines have many advantages over traditional pairwise comparisons that are suitable for longitudinal data analysis with many time points, especially for data with non-linear and complex temporal gene expression dynamics. Compared to the longitudinal transcriptomic studies from other publications with 3-5 time points where pairwise comparisons can be an effective way to reveal genes differentially expressed between time points, the study used in this pipeline contains dozens of time points, where comparing many pairs of time points can become a tedious task that also does not utilize the time-series structure fully. Moreover, it is possible to use splines to model time-course RNA-seq data with missing time points, as well as time points that are unevenly spaced, as demonstrated in Supplementary Result 4. Although there are some benefits of using splines to interpolate transcriptomic time-series data, there is a need for caution, especially when estimating parameters. When using splines to model time-course transcriptomic data, it is important to choose an appropriate number of knots, or degrees of freedom. In the user manual for limma and in Svensson’s walkthrough for sleuth on modeling time-course experiments with splines, the recommended degrees of freedom is given as 3 to 5 (25, 26). Though the reasons for this seemingly arbitrary range are not presented anywhere, this range of degrees of freedom is also recommended and seems to work well for modeling biological phenomena other than transcriptomics, such as serum glucose concentration over 30 days (10). The degrees of freedom chosen for the current pipeline was 5, to account for the fact that there are 20 and 30 data points in the two time series of interest. To evaluate the effect of degrees of freedom of the spline on the differential expression results, the current pipeline was run with degrees of freedom ranging from 3 to 7 (Supplementary Result 5).

The fit of the splines was evaluated using R^2^, with average values of 0.366 for pre-vaccination hourly samples, 0.489 for post-vaccination hourly samples, and 0.476 for daily samples. While these R^2^ values are not particularly high, several factors should be considered. Increasing the degrees of freedom generally improves the R^2^ value; however, AICc indicates that a model with five degrees of freedom is preferable to one with seven. Additionally, many time points have zero counts, even with a stricter filter than the default in sleuth, which likely contributes to the lower fit. Alternative spline approaches, such as manually placing knots, may enhance the fit, although our attempts with B-splines resulted in poor-quality genes (primarily mitochondrial), suggesting that natural splines, which are constrained to better model biological patterns, would be more suitable for this type of analysis.

## Methods

### Subsampling Time Points

We performed spline regression using sleuth (22), a program to analyze RNA-Seq experiments for which transcript abundances have been quantified with kallisto (21). Natural cubic splines were fit to the hourly series, with 20 pre-vaccination and 20 post-vaccination samples, and daily series, with 30 post-vaccination time points. Regarding sample selection for each time series, the original study selected 24 hourly samples, whereas the current analysis used only 20. We omitted the first four samples because the vaccine was administered between hours 4 and 5 of the 24-hour post-vaccination series. This omission ensured that the remaining 20 samples and their matched pre-vaccination counterparts more accurately represented the change in condition.

### Pseudo-alignment with kallisto

All samples in the time series data were pseudo-aligned to Ensembl v. 112 annotation for Homo Sapiens (GRCh38.p14) with kallisto. The k-mer length was set to the default value of 31. The `-b` argument indicates that the number of bootstraps was set to 100. Run information for each sample is assembled in Supplementary Result 6.

### Differential Expression Analysis with sleuth

All analyses were undertaken with R version 4.4.2-2024-11-01 (27) and sleuth version 0.30.1. The following R packages were loaded into RStudio: sleuth, splines, and biomaRt. The sample-to-covariates matrix, which is a required argument for sleuth, was first constructed for each of the time series. A transcript-to-gene mapping was created using biomaRt’s useEnsembl() function, in order to enable p-value aggregation of the transcript-level results (15). The annotation used for the transcript-to-gene mapping was the same annotation used for pseudo-alignment in kallisto. When importing kallisto results into sleuth, the default filter for sleuth (minimum of 5 reads observed in 47% of reads) was modified to be stricter (minimum of 5 reads observed in 67% of reads).

The objective of our study was to find genes that are highly variable in expression over time, either within one experimental condition (daily) or between two different experimental conditions (hourly) such as treatment and control, in which time and condition can interact and introduce complexities in the data. The models for the two datasets were set up as follows. For the hourly series with pre- and post-vaccination, the expression for the full model is given by:

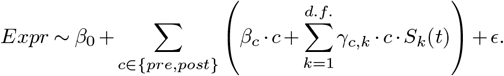

For the reduced model, the expression is given by:

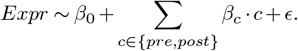

The expression weight, *β*_0_ represents the intercept term, *β*_*c*_ represents coefficients for the main effects of each condition (pre or post), *c* indicates condition (pre or post), *γ*_*c*,*k*_ represent coefficients for the interaction of condition *c* (pre or post) with the *k*-th spline basis function, *S*_*k*_(*t*) represents spline basis functions from *k* = 1 to *n* degrees of freedom, which is equal to 5 in this case, and lastly, *ϵ* is a noise term.

In the syntax of the R language, the full model for the hourly series is given by `(∼0 + group + group:X)` and the reduced model `(∼0 + group)`, where `group` is the condition covariate releveled by the pre-vaccination levels and `X` is a natural cubic spline with 5 degrees of freedom. The `(group:X)` denotes that a spline was fit to each condition, pre- and post-vaccination. The reduced model includes `group`, such that the difference between the full and reduced models represents the interaction terms, which is given as `(group:X)`. By comparing the full and reduced model in this manner, sleuth identifies isoforms that have different expression patterns for each condition. In contrast, the original study treated pre-vaccination samples as baseline levels and subtracted them from post-vaccination samples to create a ‘delta hourly’ time series for the main analysis (16). For the daily series where all time points are post-vaccination, the full model for the daily series includes time, modeled as a natural cubic spline with 5 degrees of freedom and expressed as `(∼X)` in R, and the reduced model includes noise, expressed as `(∼1)` in R. The expressions for the full and reduced model are given by:

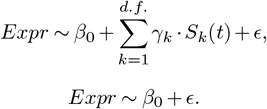

Similar to the hourly series, 0 represents the intercept term, *γ*_*k*_ represent coefficients for the *k*-th spline basis function, *S*_*k*_(*t*) represents spline basis functions from *k* = 1 to *n* degrees of freedom, which is equal to 5 in this case, and *ϵ* represents residual error.

The sleuth differential analysis tool was run on transcript mode such that splines were fit to each isoform, after which the results were compiled into gene-level differential expression results through p-value aggregation with the Lancaster method (15). There are two testing methods in sleuth: the likelihood ratio test and the Wald test. For the likelihood ratio test, which is used in the current study, sleuth compares two models: a full model with all parameters of the experiment, and a reduced model with the parameters to control for. For each type of analysis, full and reduced models were fit to the data, with specific details about the model used for each analysis provided in the Results section. To implement splines in R, the `ns()` package in the splines library was used (27). The DE analysis results were filtered for q-values < 0.05, and were exported to a Python notebook for plotting.

### Gene Set Enrichment Analysis (GSEA)

The DE analysis results from sleuth were exported as CSV files, and cleaned up in a Python notebook. The list of DE genes were analyzed with Enrichr (28) for Reactome 2024 pathways. To match the original study where the data was published, GSEA results were filtered for adjusted p-values < 0.003.

### Visualization

To visualize the spline regression results, normalized transcript counts and full models containing spline coefficients were exported as CSV files from sleuth into a Python notebook. Splines were visualized for each isoform of a DE gene, as sleuth was operated on transcript mode and the spline coefficients were fit to each isoform instead of the gene. Transcript TPMs were aggregated to plot gene-level expression.

## Supporting information

Supplementary Result 5

Supplementary Result 6

Supplementary Result 1

Supplementary Result 2

## Code and Data Availability

The data used for this study is publicly available at NCBI Gene Expression Omnibus database (accession GSE108664). The code needed to reproduce the results in this paper are available at https://github.com/pachterlab/KP_2025.

## Supplementary Note 1: Supplementary Materials

Supplementary Figures 1–4 and Supplementary Tables 1–2 are provided in the Supplementary Data. Supplementary Result 1 (a list of differentially expressed genes from the hourly and daily series), 2 (gene set enrichment analysis results), 5 (regression results using splines with alternative degrees of freedom), and 6 (details of all kallisto runs) are provided as separate .xlsx files. Supplementary Result 3 (benchmarking of our approach against edgeR-limma) and 4 (a script for running the time-course analysis pipeline on longitudinal data with uneven or missing time points) are included in the Supplementary Data.

