## Supplementary Data for "Differential Analysis Reveals Isoform Switching Following Pneumococcal Vaccination"

May 22, 2025

Supplementary Data included the following document are:

- Supplementary Figures (1–4) and Supplementary Tables (1–2)
- Supplementary Results 3 (benchmarking of our approach against edgeR-limma)
- Supplementary Results 4 (a script for running the time-course analysis pipeline on longitudinal data with uneven or missing time points)

Supplementary Results 1 (a list of differentially expressed genes from the hourly and daily series), 2 (gene set enrichment analysis results), 5 (regression results using splines with alternative degrees of freedom), and 6 (details of all kallisto runs) are provided as separate .xlsx files.

### Supplementary Figures

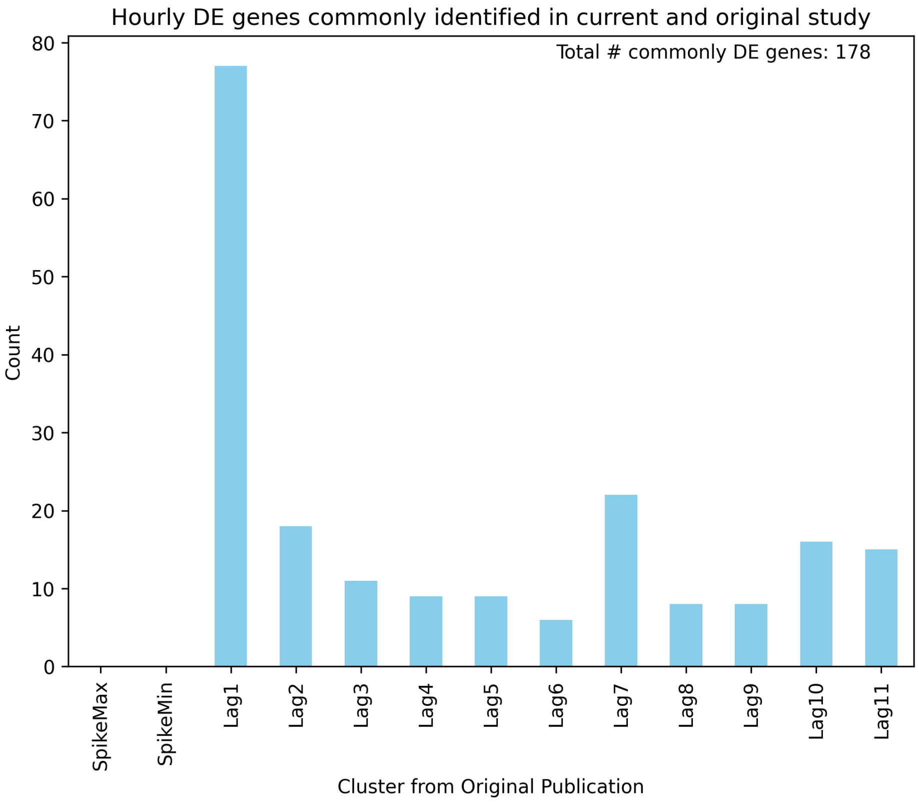


Supplementary Figure 1: Genes overlapping between sleuth's spline regression and Mias et. al.'s original study, for the hourly series. 178 of the 1208 DE genes from running sleuth on the hourly series were also found in the original study. Most of the overlapping genes were grouped into the Lag 1 cluster, which is known for providing key information about time series changes in autocorrelation methods.

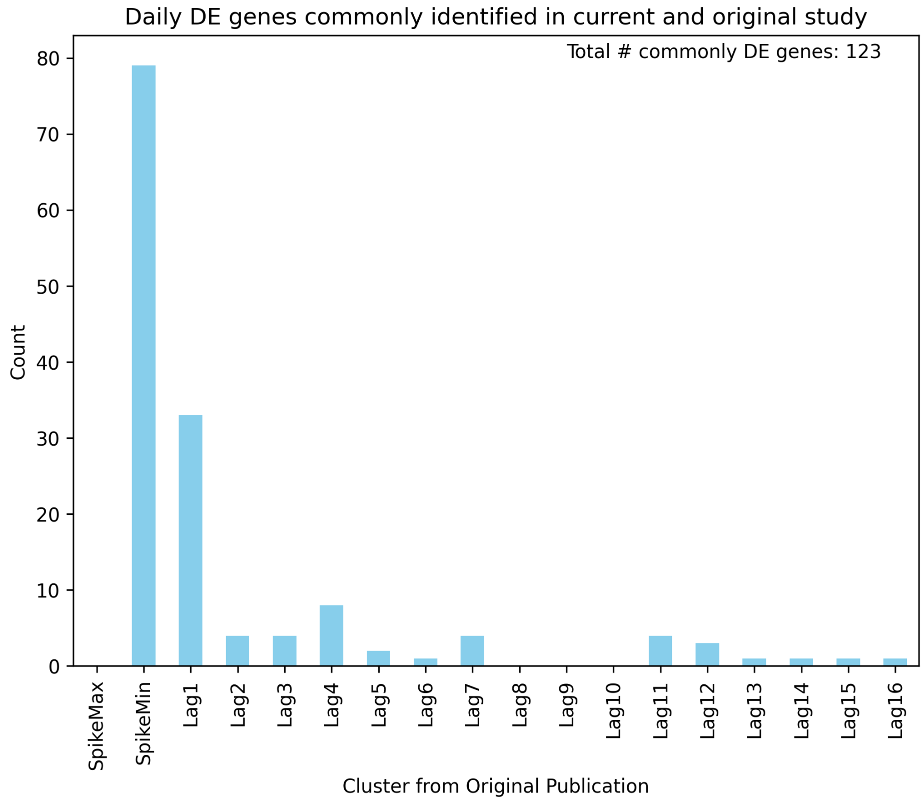


Supplementary Figure 2: Genes overlapping between sleuth's spline regression and Mias et. al.'s original study, for the daily series. 123 out of the 241 DE genes identified from running sleuth on the daily series were also marked as DE in the original study. Of the overlapping genes, most were grouped into the SpikeMin cluster, and the remainder primarily in the Lag 1 cluster.

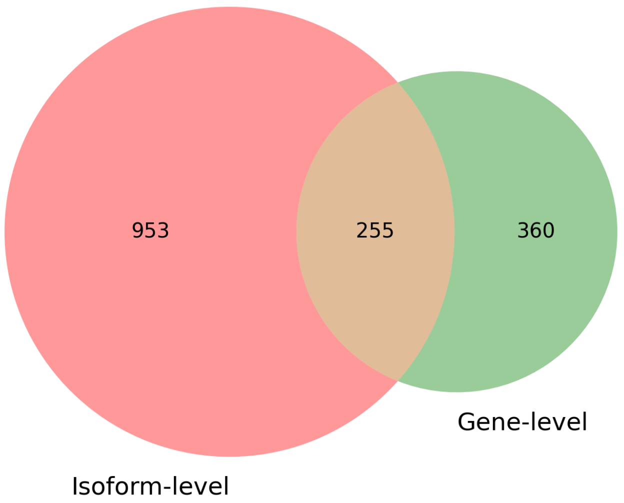


Supplementary Figure 3: Genes overlapping between isoform-level spline regression with p-value aggregation and gene-level spline regression, for the hourly series. Of the 615 genes identified in gene mode, 255 were also DE in isoform mode, but a majority of DE genes from isoform-level analysis were not DE in gene-level analysis.

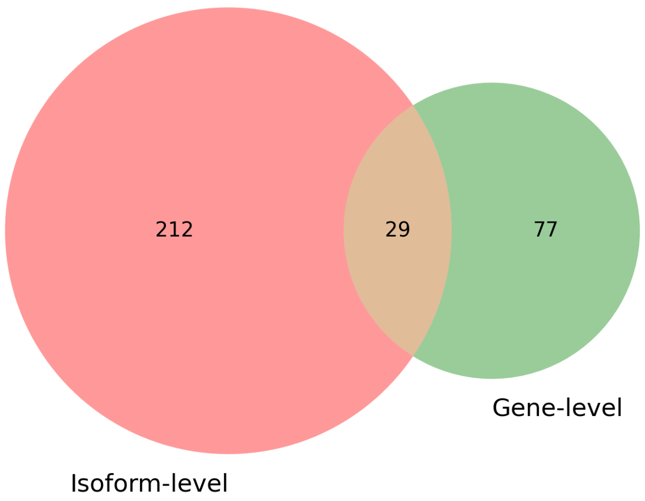


Supplementary Figure 4: Genes overlapping between isoform-level spline regression with p-value aggregation and gene-level spline regression, for the daily series. 106 genes were marked as DE in the gene-level analysis. Only 29 of the 241 DE genes from isoform-level analysis were also DE in gene-level analysis.

### Supplementary Tables

| Term | Overlap | P-value | Adj. P-value |
|---|---|---|---|
| Aerobic Respiration and Respiratory Electron Transport | 39 | 3.34E-17 | 3.85E-14 |
| FASTK Family Proteins Regulate Processing and Stability of Mitochondrial RNAs | 13 | 2.17E-16 | 8.49E-14 |
| Respiratory Electron Transport | 30 | 2.21E-16 | 8.49E-14 |
| Mitochondrial RNA Degradation | 13 | 3.55E-14 | 1.02E-11 |
| rRNA Processing in the Mitochondrion | 13 | 2.61E-11 | 6.01E-09 |
| tRNA Processing in the Mitochondrion | 13 | 1.11E-10 | 2.12E-08 |
| Complex I Biogenesis | 12 | 5.42E-07 | 8.92E-05 |
| Formation of ATP by Chemiosmotic Coupling | 7 | 9.50E-07 | 1.37E-04 |
| rRNA Processing | 22 | 1.82E-06 | 2.23E-04 |
| Mitochondrial Protein Degradation | 13 | 4.67E-06 | 5.37E-04 |

Supplementary Table 1: Top Reactome 2024 pathways from GSEA with gene-level DE genes from the hourly series. GSEA with the 615 DE genes from gene-level analysis identified Reactome pathways not particularly relevant to the immune response.

| Term | Overlap | P-value | Adj. P-value |
|---|---|---|---|
| Formation of a Pool of Free 40S Subunits | 7 | 1.34E-06 | 2.91E-04 |
| L13a-mediated Translational Silencing of Ceruloplasmin Expression | 7 | 2.46E-06 | 2.91E-04 |
| GTP Hydrolysis and Joining of the 60S Ribosomal Subunit | 7 | 2.60E-06 | 2.91E-04 |
| SRP-dependent Cotranslational Protein Targeting to Membrane | 7 | 2.91E-06 | 2.91E-04 |
| Cap-dependent Translation Initiation | 7 | 3.83E-06 | 2.91E-04 |
| Eukaryotic Translation Initiation | 7 | 3.83E-06 | 2.91E-04 |
| Formation of the Ternary Complex, and Subsequently, the 43S Complex | 5 | 8.12E-06 | 4.63E-04 |
| Peptide Chain Elongation | 6 | 9.61E-06 | 4.63E-04 |
| Eukaryotic Translation Termination | 6 | 1.22E-05 | 4.63E-04 |
| Selenocysteine Synthesis | 6 | 1.22E-05 | 4.63E-04 |

Supplementary Table 2: Top Reactome 2024 pathways from GSEA with gene-level DE genes from the daily series. GSEA with the 106 genes from gene-level analysis identified Reactome pathways that were not specific to an immune response.

### Supplementary Result 3: Comparison of pipeline performance for longitudinal RNA-seq data

The development of computational pipelines for RNA-seq analysis has significantly advanced our understanding of transcriptomic dynamics in various biological contexts. The pipeline introduced in this work using sleuth(1) is designed to perform differential expression analysis on time course RNA-seq data on the isoform level(2), which can uncover isoform-level signals that may be hidden in a traditional gene-level analysis. We applied sleuth on longitudinal, time-course RNA-seq data to see whether modeling data with splines could uncover temporal dynamics in the isoform-level, and we show that sleuth indeed was able to detect temporal signals in gene expression.

This supplementary result presents a focused comparison between our sleuth-based approach and a leading alternative approach combining edgeR for import, organization, filtering, and normalization with limma(3) for linear modeling and empirical Bayes moderation in differential expression analysis. The edgeR(4)-limma pipeline represents the most competitive alternative to our sleuth-based methodology for several key reasons. First, edgeR provides a direct method for importing kallisto(5) pseudo-alignment output through tximport(6), allowing us to use identical isoform-level quantification data as input to both pipelines. This enables a more objective comparison by eliminating variability that might arise from different alignment strategies. Second, limma explicitly supports spline regression—the same statistical approach employed in sleuth—in their user guide, making it theoretically capable of modeling similar expression patterns across time. The comparison is focused on the hourly time series, which has 20 pre-vaccination and 20 post-vaccination time points, to highlight the viability of using these computational pipelines on time series data that have matched time points in different conditions.

Despite these methodological similarities, our preliminary analyses revealed substantial differences in the ability of these pipelines to detect biologically meaningful signals in longitudinal RNA-seq data. While both approaches utilize comparable statistical frameworks, our sleuth-based pipeline demonstrated enhanced sensitivity in identifying time-dependent expression patterns with biological relevance. By benchmarking against the edgeR-limma approach—which represents a widely-used practice in the field—we provide evidence for the specific advantages of our sleuth-based pipeline in the context of longitudinal RNA-seq analysis. The results presented in this supplementary serve to validate the technical advancements of our approach and provide practical guidance for researchers analyzing longitudinal transcriptomic data who must choose between these competing methodological frameworks.

```r
# Import dependencies
library(tximport, tidyr)
library(limma)
library(splines)

# Set working directory and import metadata
setwd("~../../longsaliva")
s2c <- read.delim("s2c_hourly.txt", sep=" ",
                  header=TRUE)
df <- dir(file.path("."))
time <- rep(seq(from=1, to=length(s2c$sample)/2, by=1),
            times=2)
s2c <- dplyr::mutate(s2c, time=time)
colnames(s2c) <- c("path", "sample", "condition",
                   "time")

# Set up annotations for genes and transcripts
ensembl <- biomaRt::useEnsembl(biomart = "genes",
                       dataset = "hsapiens_gene_ensembl",
                       mirror = "useast")
t2g <- biomaRt::getBM(attributes = c("ensembl_transcript_id",
                                      "ensembl_gene_id"),
                      mart = ensembl)
t2g <- dplyr::rename(t2g, TXNAME = ensembl_transcript_id,
                     GENEID = ensembl_gene_id)
files <- file.path(s2c$path, "abundance.h5")
names(files) <- s2c$sample
all(file.exists(files))
```

```r

# Import kallisto outputs
txi <- tximport(files, type = "kallisto",
                tx2gene = t2g,
                ignoreTxVersion = TRUE,
                countsFromAbundance = "lengthScaledTPM")
names(txi)

# Filter low-expression counts with edgeR
y <- edgeR::DGEList(txi$counts)
keep <- edgeR::filterByExpr(y)
y <- y[keep, ]

# Modeling time as natural spline with 5 d.f. as with sleuth
X <- ns(s2c$time, df=5)

#Then fit separate curves for the control and treatment groups:
Group <- factor(s2c$condition)
Group <- factor(s2c$condition,
                levels = c("pre_vaccination", "post_vaccination"))
design <- model.matrix(~0 + Group*X, data=y)
colnames(design)

# Normalize and run voom transformation on limma
y <- edgeR::calcNormFactors(y)
v <- voom(y, design)
fit <- lmFit(v, design)
fit <- eBayes(fit)

# View DE results
tt <- topTable(fit, coef=8:12)
```

The edgeR-limma pipeline above yielded only 10 DE genes, which was not a sufficient number of genes to yield any significant gene set enrichment analysis (GSEA) results. The genes marked as differentially expressed over time are shown below:

| Gene ID | Gene Name | Avg. Expr. | F-statistic | P.Value | Adj.P.Val |
|---|---|---|---|---|---|
| ENSG00000274058 | None | 4.783 | 11.286 | 6.20E-07 | 4.43E-03 |
| ENSG00000135390 | ATP5MC2 | 3.067 | 10.923 | 8.98E-07 | 4.43E-03 |
| ENSG00000101146 | RAE1 | 1.053 | 9.718 | 3.23E-06 | 1.06E-02 |
| ENSG00000006534 | ALDH3B1 | 2.288 | 9.375 | 4.70E-06 | 1.16E-02 |
| ENSG00000149573 | MPZL2 | 2.284 | 9.027 | 6.96E-06 | 1.21E-02 |
| ENSG00000168421 | RHOH | 5.018 | 8.879 | 8.22E-06 | 1.21E-02 |
| ENSG00000083844 | ZNF264 | 2.680 | 8.845 | 8.55E-06 | 1.21E-02 |
| ENSG00000141665 | FBXO15 | 2.431 | 8.566 | 1.18E-05 | 1.36E-02 |
| ENSG00000189266 | PNRC2 | 3.531 | 8.520 | 1.24E-05 | 1.36E-02 |
| ENSG00000151366 | NDUFC2 | 2.267 | 8.263 | 1.68E-05 | 1.49E-02 |

Supplementary Table 3: Genes identified as differentially-expressed over time by the edgeR-limma pipeline on the hourly time series. Only 10 genes were identified, as opposed to the 1208 identified by sleuth on the same data. Gene Set Enrichment Analysis was not performed due to the insufficient number of genes.

The comparative analysis between our sleuth-based pipeline and the edgeR-limma approach revealed striking differences in sensitivity for detecting differential expression in longitudinal RNA-seq data. The edgeR-limma pipeline yielded only 10 differentially expressed genes, despite using identical isoform-level quantifications from kallisto and employing comparable statistical frameworks for modeling temporal patterns. Our sleuth-based methodology identified 1,208 differentially expressed genes from the same dataset, demonstrating significantly higher detection power.

The substantial disparity in detection sensitivity had significant downstream implications for biological interpretation. The limited set of 10 genes identified by edgeR-limma proved insufficient to yield any statistically significant results in gene set enrichment analysis (GSEA), effectively preventing meaningful pathway-level insights. In contrast, the much larger gene set identified by sleuth enabled robust enrichment analysis, revealing biologically coherent temporal patterns as detailed in the main text.

These results underscore the critical importance of pipeline selection in longitudinal RNA-seq studies, particularly for isoform-level analyses. While both methods theoretically support spline-based modeling of time-course data, sleuth demonstrated dramatically enhanced capacity to detect biologically relevant expression changes. This performance difference may be attributed to sleuth's specialized design for transcript-level analysis and its sophisticated handling of technical variance in RNA-seq data.

For researchers conducting longitudinal transcriptomic studies, these findings suggest that sleuth offers substantial advantages over the edgeR-limma approach, particularly when temporal patterns and isoform-specific dynamics are of interest. The practical implications extend beyond mere statistical power to the fundamental ability to extract meaningful biological narratives from complex time-series RNA-seq data.

### Supplementary Result 4: Missing and uneven time points in longitudinal data

The following code can be used for longitudinal data with missing or unevenly-distributed time points. We demonstrate that our analysis pipeline can detect genes differentially expressed over time between two experimental conditions, by sampling time points from the time series data we used. Even when the time points between the experimental conditions are not matched, this pipeline still works.

```r
library(sleuth, splines)
library(biomaRt)
ensembl <- biomaRt::useEnsembl(biomart = "genes",
                               dataset = "hsapiens_gene_ensembl",
                               mirror='useast')
t2g <- biomaRt::getBM(attributes = c("ensembl_transcript_id",
                                     "ensembl_gene_id",
                                     "external_gene_name"),
                      mart = ensembl)
t2g <- dplyr::rename(t2g, target_id = ensembl_transcript_id,
                     ens_gene = ensembl_gene_id,
                     ext_gene = external_gene_name)

setwd("~/longsaliva")
s2c <- read.delim("s2c_hourly.txt", sep=" ", header=TRUE)
time <- rep(seq(from=1, to=length(s2c$sample)/2, by=1), times=2)
s2c <- dplyr::mutate(s2c, time=time)
sample <- paste0(rep(c('pre', 'post'), each=20),'_',s2c$time)
s2c$sample <- sample

colnames(s2c) <- c("path", "sample", "condition", "time")
# If the time points between pre- and post- are matched
# s2c <- s2c[c(1, 2, 4, 8, 20, 21, 22, 24, 28, 40), ]

# This works when time points aren't matched too
s2c <- s2c[c(1, 2, 4, 8, 20, 21, 23, 26, 31, 40), ]

group <- relevel(factor(s2c$condition), ref='pre_vaccination')

new_filter <- function(row, min_reads = 5, min_prop = 0.67) {
  mean(row >= min_reads) >= min_prop
}

so <- sleuth_prep(s2c, target_mapping = t2g,
                  aggregation_column = "ens_gene",
                  extra_bootstrap_summary = TRUE,
                  filter_fun = new_filter)

X <- splines::ns(s2c$time, df=5)
full_design <- model.matrix(formula(~0 + group + group:X))
colnames(full_design)

so <- sleuth_fit(so, full_design, "full")
so <- sleuth_fit(so, ~0+group, "reduced")
so <- sleuth_lrt(so, "reduced", "full")

sleuth_table <- sleuth_results(so, 'reduced:full', 'lrt', show_all = FALSE, pval_aggregate =
     TRUE)
sleuth_de <- dplyr::filter(sleuth_table, qval <= 0.05)
head(sleuth_de)
```

[6] Charlotte Soneson, Michael I Love, and Mark D Robinson. Differential analyses for RNA-seq: transcript-level estimates improve gene-level inferences. 4:1521. doi: 10.12688/f1000research.7563.2. URL `http://dx.doi.org/10.12688/f1000research.7563.2`.
